# Phylogeographic analysis and identification of factors impacting the diffusion of Foot-and-Mouth disease virus in Africa

**DOI:** 10.1101/358044

**Authors:** Florian Duchatel, Mark Bronsvoort, Samantha Lycett

## Abstract

Foot and mouth disease (FMD) is endemic in sub-Saharan Africa and can lead to important and continuous economic losses for affected countries. Due to the complexity of the disease epidemiology and the lack of data there is a need to use inferential computational approaches to fill the gaps in our understanding of the circulation of FMD virus on this continent. Using a phylogeographic approach we reconstructed the circulation of FMD virus serotypes A, O and SAT2 in Africa and evaluated the influence of potential environmental and anthropological predictors of virus diffusion. Our results show that over the last hundred year the continental circulation of the tree serotypes was mainly driven by livestock trade. Whilst our analyses show that the serotypes A and O were introduced in Africa trough livestock trades, the SAT2 serotype probably originates from African wildlife population. The circulation of serotype O in eastern Africa is impacted by both indirect transmission through persistence in the environment and anthropological activities such as cattle movements.

---

Foot and mouth disease (FMD) affects more than 70 species of cloven-hoofed animals ^1^. The disease is characterised by the development of vesicles in and around the mouth, on the feet and other sites of the skin ^1^. The causal agent is a positive-sense, single-stranded RNA virus of the *Picornaviridae* family ^2^ called foot-and-mouth disease virus (FMDV). Its genome encodes the information for 4 structural proteins (VP1-4) and 8 non-structural proteins (7 proteases and one RNA polymerase). Based on the level of cross protection between strains, the virus can be divided into seven serotypes, O, A, C, Southern African Territories [SAT] 1, SAT 2, SAT 3 and Asia 1 ^3,4^, which are clinically indistinguishable from each other but which have different epidemiologies. The hosts that are considered to play an active role in these epidemiologies are cattle, buffaloes, pigs, sheep and goats ^5^.

FMD susceptibility varies according to the host and strain of FMDV involved. The severity of the infection depends of the amount of virus inoculated, the serotype, the host species and the individual immunity ^6^. The commonest route of infection for a new host is by direct contact with an infected animal ^1,7^. The infection may also occur indirectly by contacts with contaminated surfaces or products, such as infected personnel, vehicle or fomites ^5^. Movement of animals and animal products are considered to play an important role in the disease circulation in endemic areas and are considered the main factors for FMDV transboundary spread ^8^.

FMD has been eradicated in many high income countries but is still endemic in numerous low and middle income countries (LMICs) ^9^ particularly in Africa and South and East Asia. Although FMD has a low mortality rate in adult animals it causes significant productivity losses that may lead to important and continuous economic losses for farmers and impact countries trading ability at a national level ^8^. Although work has been done to understand the impact of FMDV in large scale dairy farms in LMICs ^10^ there is still a lack of data to quantify its impact more broadly on the economy of endemically infected countries ^11^.

FMDV is endemic in most of sub-Saharan Africa with an epidemiology considered to be more complex than in other regions of the world due to multiple serotype and wildlife reservoirs ^12^. However, due to a general lack of surveillance and animal traceability, very few statistics on disease incidence and circulation exist for Africa. Although there are a few studies on animal trade and seasonal migration of nomadic and pastoralist herds in sub-Saharan African ^8,–12,14^ we need analytical approaches that use existing data to improve our understanding of the circulation of FMDVs in this part of the world.

Many wildlife species can be infected by FMDVs in Africa ^6^ but amongst all these potential hosts, only the Cape buffalo (*Syncerus caffer*) and impala have been implicated in the transmission of FMDV to domestic cattle ^15^. Even though the Cape buffalo is suspected to be the primary reservoir and the main source of SAT serotypes in Southern Africa^16^, its role as a viral source for livestock epidemics for the FMDV O, A and C serotypes elsewhere in Africa is still unclear^11,13,17,18^, and might be inexistent^19^.

It has already been observed that the spatiotemporal occurrence and circulation of FMDV in Africa is mainly affected by human activities though domestic animal movements^20–23^. However, several environmental characteristics and attributes such as the landscape, vegetation, natural barriers to animal movements such as roads, rivers or mountains have the potential to influence the dynamics and circulation of FMD ^24–26^.

Because FMDV are single stranded RNA viruses and lack a proof-reading mechanism for their genome replication they are subject to a high evolutionary mutation rate^4^. The history of these mutations can provide information on the ecological processes and population events that shaped the virus evolution even if not directly observed. These processes, along with other evolutionary parameters, can be modelled while reconstructing the phylogenetic trees^27,28^. Furthermore, by combining genetic data and spatial information, phylogeographic tree reconstruction can be used to estimate the unobserved geographical circulation of a pathogen^29^. Virus movements can be modelled as discrete transmission events between the sampled locations^30^ or as a continuous process using different random walk diffusion models^31^. Recently both discrete and continuous approaches have been extended to test and quantify the contribution of potential environmental and anthropological parameters (predictors of viral diffusion) that might influence the spread and circulation of the studied pathogen ^32,33^.

The aim of this paper is to gain a better understanding of the circulation of FMDV in Africa, comparing discrete and continuous approaches^32,34^. A detailed discrete phylogeographical analysis of serotypes A, O and SAT2 sequences was performed, and the influence of 13 potential environmental and anthropological predictors of virus diffusion were quantified and tested using both discrete and continuous approaches.

## RESULTS

### Discrete phylogenetic analysis

Evolutionary parameters estimation

Using a relaxed log-normal clock and a skygrid population model for the serotype A and O; and a constant clock and skygrid population model for the serotype SAT2 we estimated evolutionary parameters and reconstructed the phylogenetic trees (see Figure 1). Overall, we observed a mean mutation rate of 4.67×10^−3^ substitutions per site per year and 3.69×10^−3^ for the serotypes A and O respectively. We also estimated a significantly slower mutation rate of 1.1×10^−3^ for the serotype SAT2 (see supp. table 1).

**Table 1:**
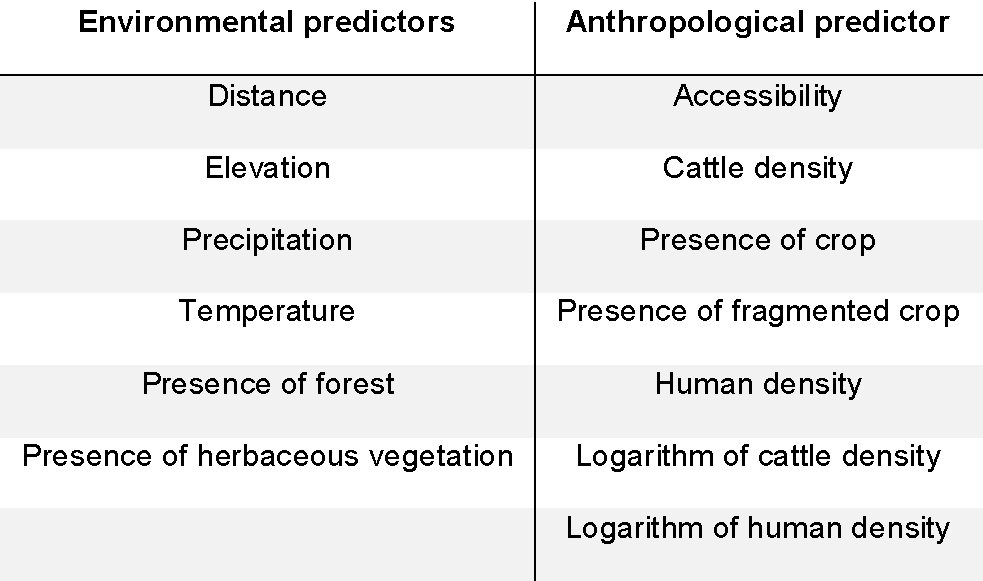
Environmental and anthropological predictors tested for an effect on the FMDV serotype O diffusion in Eastern Africa

**Figure 1:**
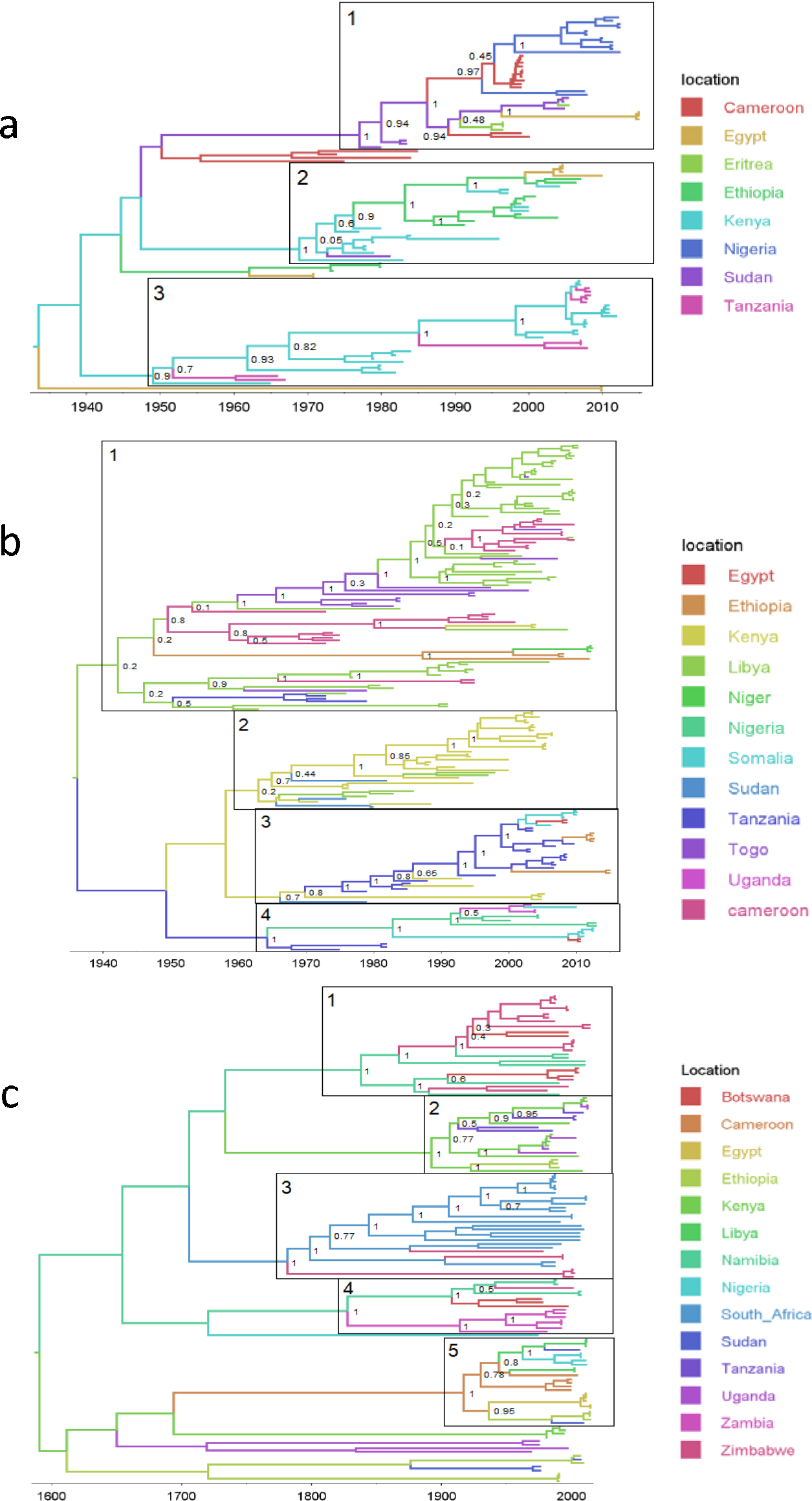
Bayesian MCC time scaled discrete phylogeographic tree for the three studied serotypes. a. Bayesian phylogeographic tree for serotype A using 107 VP1 sequences. b. Bayesian phylogeographic tree for serotype O using 192 VP1 sequences. c. Bayesian phylogeographic tree for serotype SAT2 using 135 VP1 sequences. The phylogeny branches are coloured according to their descendent nodes location with the key for colours shown on the right. The main clades for each of the studied serotypes are identified on the phylogeographic trees. The nodes of the isolated clades are annotated with their posterior probabilities.

### Phylogeographic tree reconstruction for serotype A

The reconstructed phylogeographic tree of the African serotype A viruses has a time to most recent common ancestor (TMRCA) of around 1926 (1889.6 – 1950 95%HPD), with geographic origin in the eastern part of Africa and high posteriors probabilities for Kenya (49.83 %) and Ethiopia (35.95 %) (see fig. 1a). For serotype A there is no clear clade separation between the western and eastern sides of Africa, as the first isolated clade combines all the western African sequences as well as sequences from Sudan, Ethiopia and Egypt. Although a few transmissions events are observable between the two sides of Africa, all of them involve Sudan as a link between them.

### Phylogeographic tree reconstruction for serotype O

The TMRCA of the African serotype O is estimated to be 1937 (1921 – 1952 95%HPD) and located in the eastern part of Africa with high posteriors probabilities for Kenya (61.49 %), Sudan (17.15 %) and Uganda (11.42 %) (see fig.1b). The reconstructed phylogeographical tree is composed of four geographically defined main clades. The first clade is almost entirely composed of Kenyan, Tanzanian and Ugandans sequences with only a few transmissions to other countries. The second clade is mostly situated in Ethiopia with few transitions to Kenya and Somalia. The third clade is centred in Sudan with incursions into Nigeria, Cameroon, Egypt and Ethiopia. The fourth clade is centred in West and Central African countries (Cameroon, Nigeria, Niger and Togo) and seems to originate from Sudan. Overall, we can see that the situation for the serotype O is quite similar to that of serotype A with only few observed transmissions between the Eastern and western sides of Africa, with Sudan acting as a link between the two sides of Africa.

### Phylogeographic tree reconstruction for serotype SAT2

The diversity for serotype SAT2 viruses is much greater than for serotypes A and O, and the TMRCA is much older, estimated as 1583 (1722 – 1440 95% HPD). Due to these long timescales and low posterior probabilities on the location it is difficult to estimate an origin location (see fig.1c).

Five geographically defined main clades, with location posterior probabilities above 45% can be observed. The first clade is exclusively composed of sequences from Botswana, Namibia and Zimbabwe and seems to have its origin in the first half of the 19^th^ century. The second clade is composed of Ethiopian, Kenyan, Ugandan and Tanzanian sequences and seems to originate at the transition between the 19^th^ and 20^th^ century. The third clade seems to have emerged at the end of the 18^th^ century and is composed of Zimbabwean and all the South-African sequences. The fourth clade has its TMRCA in the first half of the 19^th^ century and is composed of sequences from Botswana, Namibia and Zambia. The last clade is the most more diverse and emerged over the last century and is composed of Eastern, Western and Northern African sequences (Cameroon, Egypt, Ethiopia, Libya, Nigeria and Sudan). Compared to the other serotypes, the SAT2 serotype seems to have appeared first in Southern Africa before moving to other parts of the continent; with a separation between Southern countries and the other African countries. Whilst the presence of the virus in Southern Africa seems to be geographically defined, the virus seems to circulate more freely amongst the other African countries.

### Bayesian stochastic search variable selection analysis

Using a Bayesian stochastic search variable selection (BSSVS) analysis we identified well supported rates of transition between the sampled countries. The support for the rates was quantified with Bayes factors (BF), and rates with BF >=3 are shown in Figure 2. Globally, the results for the serotypes A and O look quite similar, with Sudan acting as a link between the Eastern, Northern and Western part of the continent (see fig. 2a and 2b). For both serotypes there is a clear transmission route starting from Ethiopia, passing through Kenya to Tanzania. Although most of the observed transmission routes have low BF values, the situation for the SAT2 serotype is fairly similar to what is observed for the serotypes A and O (see fig.2c). For SAT2 multiple transitions rates can be observed within Eastern and western Africa with Sudan acting as link between the two sides. However, with only two rates linking South-Africa to the rest of the continent, Southern-African countries are quite isolated from the other African countries (see supp table 1,2 and 3).

**Table 2:** Bayes factor values associated with the effect of each predictor on the connectivity between the sampled locations using a discrete or continuous location approach. Each predictor raster was used as conductance or resistance to evaluate if the predictor have a positive or negative effect on the viral genetic connectivity.

| Predictor | Discrete locations |  | Continuous locations |  |
| --- | --- | --- | --- | --- |
|  | Conductance | Resistance | Conductance | Resistance |
|  | Bayes factor | Bayes factor | Bayes factor | Bayes factor |
| Accessibility | 0 | 8 | 0 | 0 |
| Cattle density difference | - | 0 | - | - |
| Cattle density | 2 | 0 | 0 | 3 |
| Presence of crop | 0 | 0 | 1 | 0 |
| Presence of crop (combined) | 0 | 0 | 0 | 4 |
| Distance | - | 8 | - | - |
| Elevation | 1 | 0 | 0 | 1 |
| Presence of forest | 1 | 0 | 1 | 0 |
| Presence of fragmented crop | 1 | 0 | 0 | 6 |
| Human density | 1 | 0 | 1 | 0 |
| Precipitation | 2 | 7 | 0 | 0 |
| Presence of herbaceous vegetation | 1 | 0 | 0 | 0 |
| Temperature | 4 | 7 | 0 | 1 |
| Logarithm Cattle density | 4 | 1 | 1 | 0 |
| Logarithm Human density | 9 | 2 | 0 | 0 |

**Figure 2:**
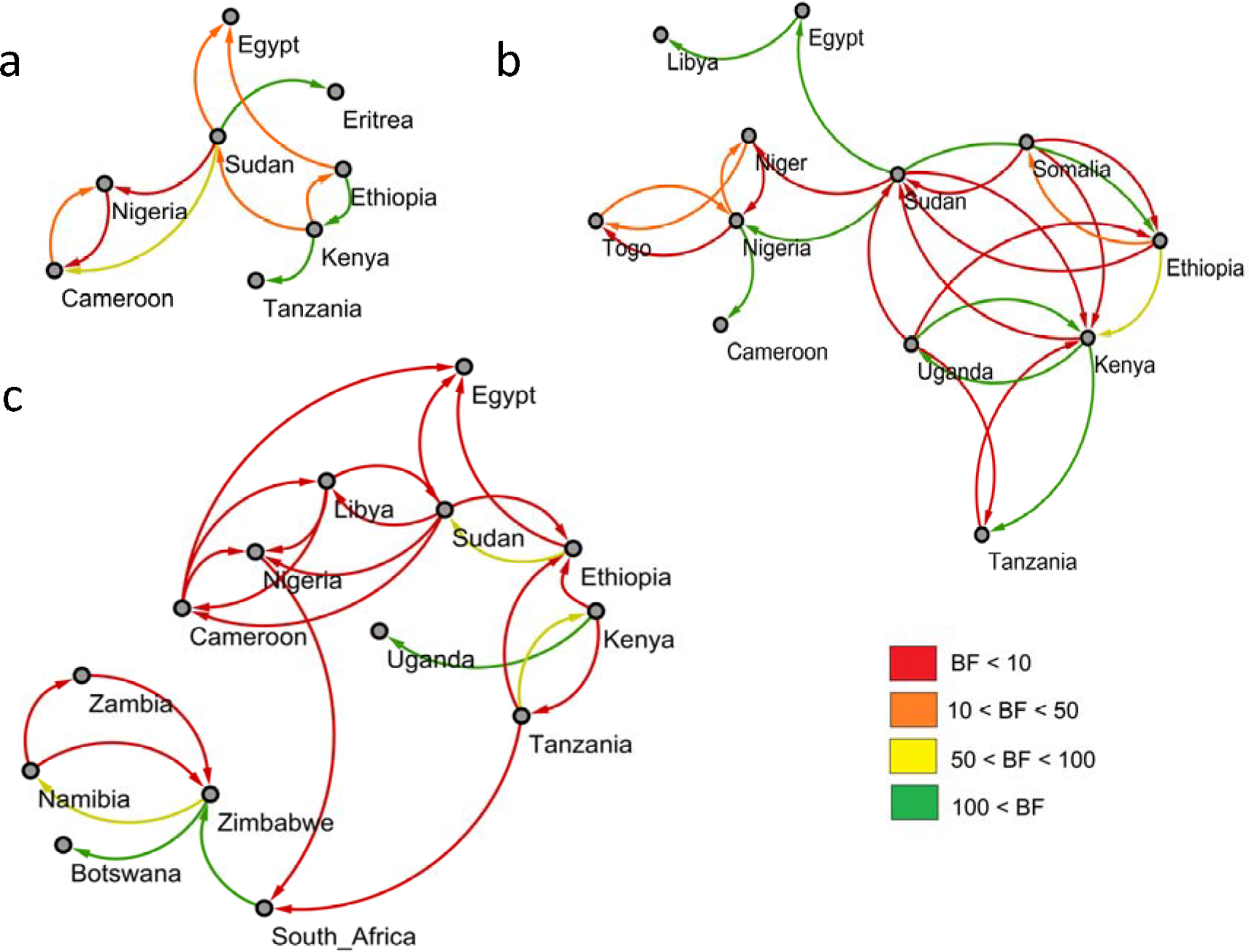
Outputs of the BSSVS analysis for the three studied FMDV serotypes showing the best supported rates of transition between the sampled countries. The edges colours represent the relative strength by which the rates are supported (red: X-Y, yellow: X-Y, green: >= Z). a. For FMDV serotype A. b. For FMDV serotype O. c. For FMDV serotype SAT2.

**Figure 3:**
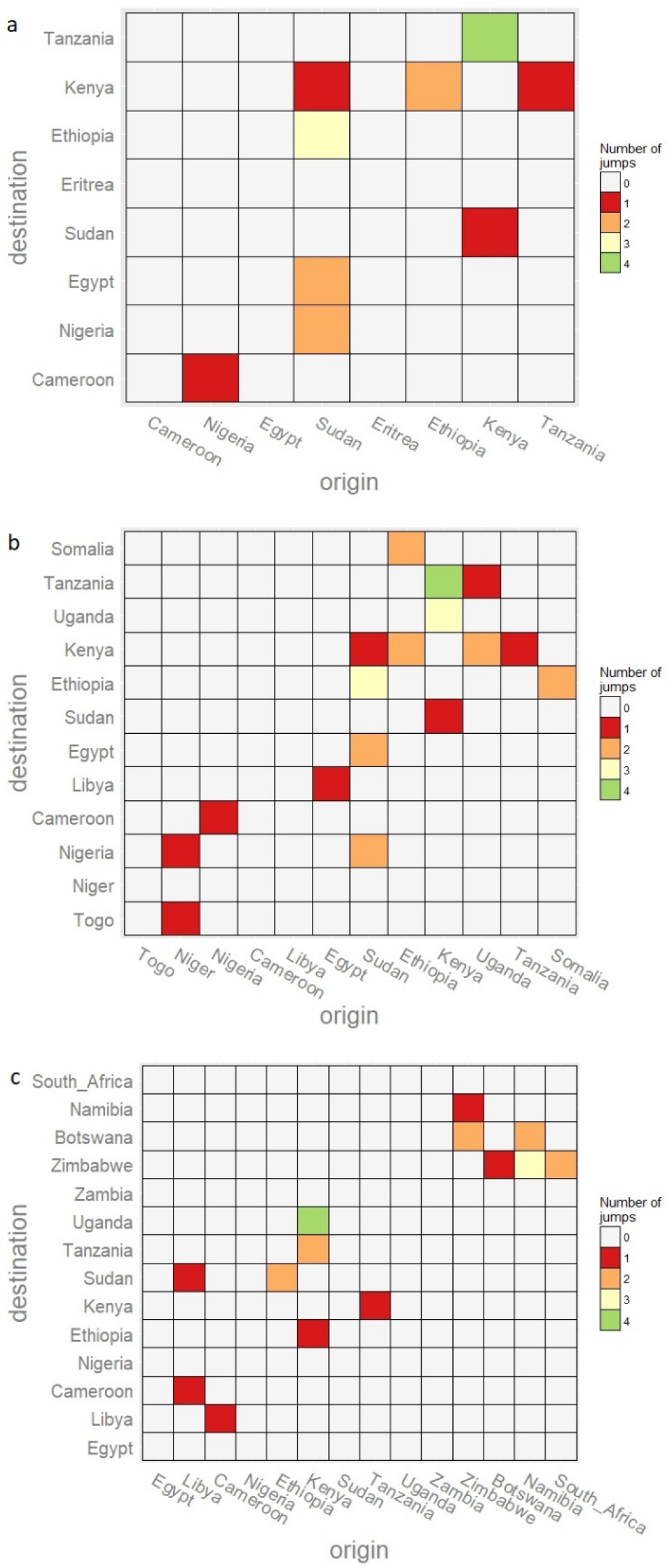
Heatmap showing the number of transitions between the sampled countries for the three studies FMDV serotypes. The heatmaps are coloured according to the number of observed transitions between countries. a. FMDV serotype A. b. FMDV serotype O. c. FMDV serotype SAT2

### Markov jumps analysis

To complement the BSSVS analysis, an estimation of the number of transmission events between the different locations using a Markov jump analysis was performed. For both FMDV serotypes A and O we observe some transmission events starting in Kenya toward other East African countries such as Tanzania, Ethiopia and Uganda. For these two serotypes we also detected jumps from Sudan in the direction of North-Eastern and Western African countries such as Egypt, Eritrea, Cameroon and Nigeria. Therefore, it seems that for these two serotypes Kenya and Sudan act as sources for the virus, but toward different directions (see supp. table 4 and 5). For the serotype SAT2 the situation is quite different. In this case we observed fewer transition events between the different sampled countries (see supp. table 6), with most of the observed transitions occurring within Eastern African and Southern Africa with no clear link between them.

### Environmental and anthropological factors affecting FMDV diffusion

Using the output from previous discrete phylogeographical analysis we isolated a FMDV serotype O monophyletic clade with a time to the most recent common ancestor (TMRCA) below 25 years and a high posterior probability on the location for all its nodes. The selected 46 sequences originated from Kenya, Tanzania and Uganda (See supp. figure 1 and supp. table 8). Using a general linear model (GLM) for the discrete location approach and the recently developed *SERAPHIM* package^34^ for the continuous location approach, we tested the impact of 13 different anthropological and environmental factors (predictors) on the FMDV diffusion in Eastern Africa.

### Predictive factors for FMDV diffusion using a discrete location approach

A generalized linear model (GLM) was used to parametrize the transition rate matrices between the sampled locations as a function of our selected predictors^32^ on a posterior set of time-resolved trees. We considered the set of predictors to be ‘conductors’ – i.e. enhancing viral diffusion, or ‘resistors’ – i.e. impeding viral diffusion. We observed that the diffusion process was enhanced by the average daily temperature (BF 4), the logarithm of the cattle density (BF 4) and human densities (BF 9) (see table 2). It was impeded by the accessibility (BF 8), the distance between sampled locations (BF 8), average amount of precipitation (BF 7) per year and by the average daily temperature (BF 7) (for all the results see supp. Table 9 and 10). To gain a better understanding of the impact of the average temperature and precipitation on the viral diffusion we selected different thresholds of precipitation and temperature to parametrize our GLM analysis (see supp. table 11 and 12). We detected that low precipitation values (< 80 mL/year) were associated with an impeding (negative) impact on the viral diffusion processes whereas high precipitation was associated with an enhancing (positive) effect on the diffusion process. We also observed that in the case of low temperature (below 22°C) a positive effect on the diffusion could be observed whereas temperatures around °C had a negative effect on virus diffusion. Temperatures above 24°C seemed again to have a positive impact on the virus spread. It was difficult to distinguish between the effects of accessibility and human density because the two were strongly negatively correlated confounding the analysis (see sup. table 13).

**Figure 4:**
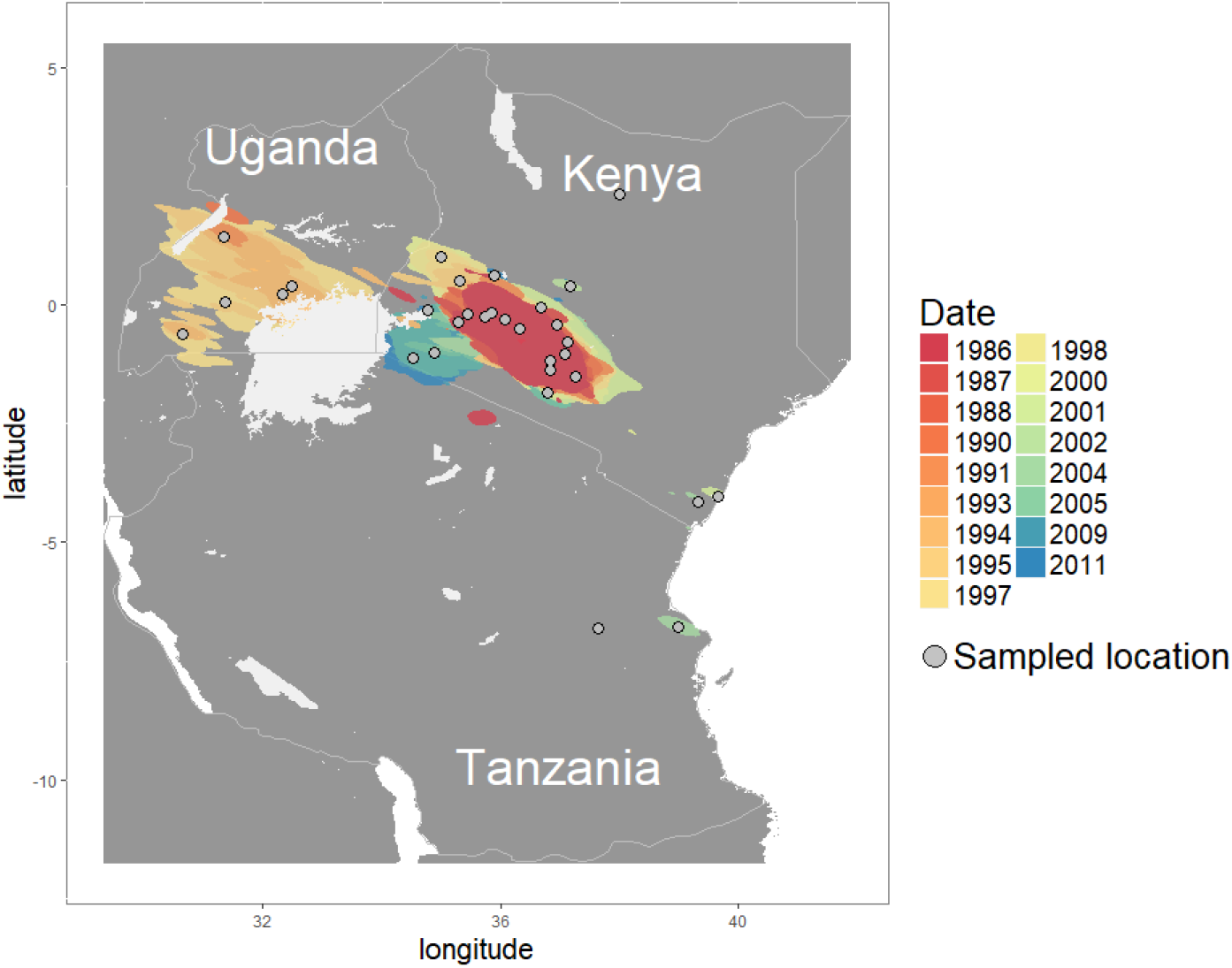
Map showing the continuous diffusion of the isolated clade of FMDV serotype O, with the sampled locations as grey circles. The virus movments were reconstructed using a random walk model with a underlying lognormal distribution

### Predictive factors for FMDV diffusion using a continuous diffusion approach

Using a random walk model, we were able to reconstruct the virus diffusion in a continuous setting for the isolated FMDV serotype O. Using the R package *SERAPHIM* ^34^ we evaluated the impact of the predictors on the virus diffusion and observed an impeding (negative) influence of the cattle density (BF 3), the presence of cropland (fragmented cropland and pure cropland areas combined) (BF4) and by the presence fragmented cropland (BF6). We were not able to detect a predictor with an enhancing (positive) influence on the diffusion process (see table 2).

To gain a better understanding of the role of the fragmented crop and cattle density we isolated the areas newly covered over the course of the infection and observed how the presence of the two predictors evolved. Overall, we noticed an opposite trend in how their densities evolved with the elapsed time (see supp.fig.2). For the fragmented crop density, high values of crop densities became more common over the course of the epidemic, with the disease moving from areas with low crop densities to areas with high crop densities. For the cattle density the opposite trend was observed with high values of cattle density more common at the start of the epidemic, the disease starting in an area where cattle densities were high and moving toward areas with lower densities of cattle. To better understand the effect that the cattle density had on virus diffusion we looked at selected areas above different thresholds of cattle density and used them as inputs in in the *SERAPHIM* package. By doing so we were able to observe that densities of cattle above 125 cattle per square km have the biggest negative impact on the virus diffusion (see supp. table 14).

## DISCUSSION

In this paper we have applied the most recent existing phylogenetic methods on all available African FMDV VP1 sequences for the serotypes A, O and SAT2. Our work has some limitations, especially regarding the limited availability of sequences. Our sampling is obviously unbalanced as it is based on submissions by individual countries or ad hoc research projects and the effect that it has on the results quality is uncertain. Increasing the number of available FMDV sequences from diverse locations and hosts would help to develop models that better represent the diffusion of FMDV in Africa and lead to better environmental and anthropological effect estimation.

The estimated substitution rates of 4.67×10^−3^, 3.69Ex10^−3^ and 1.1×10^−3^ mutations per site per year for the serotypes A, O and SAT2 from our results are similar to previous estimates of 4.26×10^−3^, 3.14×10^−3^ and 1.07×10^−3^ mutations per site per year for the same serotypes as found by Tully et al^35^. Overall, we observed similar evolutionary patterns for both FMDV serotypes A and O. Those serotypes seem to have appeared in Eastern Africa around 1930. Our results pointed to the possible role of Kenya as a viral source for East African countries and the role of Sudan as a link between East Africa and North-East Africa. The evolution of the SAT2 serotype seems to be quite different. According to our results, this serotype seems to have emerged around 1583. Over the 5 main clades that compose the SAT2 phylogeny only two emerged outside Southern Africa (clades 2 and 5). Those two clades involved Eastern (Kenya, Ethiopia and Tanzania), or Western/Central/Northern African countries (Libya, Nigeria, Cameroon and Egypt) with Sudan being the only country present in both clades. Additionally, these two clades have emerged more recently than those involving Southern African countries (second part of 19^th^ century/early 20^th^ century for the clade 2 and 5 against late 18^th^ century/early 19^th^ century for the other clades).

For the FMDV serotypes A and O the observation of well supported rates of viral transmission between Eastern Africa and Western Africa can be explained by the existence of commercial routes between those areas. It is indeed acknowledged that livestock trades play an important role in FMDV circulation in sub-Saharan Africa, through trading routes exist between the Horn of Africa and Eastern Africa, with Sudan acting as key commercial intermediate^8^. Additionally, the existence of a relatively recent common ancestor for the FMDV serotypes A and O is further support for the idea that these serotypes were imported into Africa at the start of the 19^th^ century through livestock trade from Asia and Europe^11^.

The SAT2 serotype analysis shows the signs of the impact that the African rinderpest epidemics had on FMDV circulation in Africa. Although FMDV was first reported in southern Africa in 1795 it likely had coevolved with buffalo over millennia resulting in a large diverse viral pool, but the rinderpest epidemic decimated almost all FMDV potential carriers and probably pushed it through a huge bottleneck^36^. It is thought that FMDV reemerged from wild buffalo population that survived the rinderpest epidemic, before being reported again in 1932 in Southern Africa^11^. Consistent with this hypothesis, of all the clades present in the reconstructed phylogeny, only those originating from Southern African countries have an TMRCA older than the African rinderpest epidemic (clades 1, 3 and 4). The SAT2 serotype probably spread into non-Southern African countries through infected livestock movements, increasing the virus mobility and explaining the more recent TMRCA and the highest country diversity observed in the more recent clades (clades 2 and 5). Additionally, with the observation of similar transition rates patterns amongst the three serotypes, the BSSVS analysis suggest that over the last hundred years the SAT2 circulation between African countries was mainly driven by domestic animal movements, with the relative isolation of southern Africa being the result of the different control measures in place in this region^37^.

Using both a discrete and continuous framework, we looked at the effect that diverse environmental and anthropological factors had on the diffusion of an isolated FMDV serotype O clade that circulated in Kenya, Uganda and Tanzania. The results of the discrete approaches suggest that the FMDV diffusion is facilitated by low average daily temperature (<22°C), high averages precipitations (>80 mL/year) as well as high human and cattle densities. We were also able to observe that the virus diffusion was negatively impacted by the accessibility (long travel time needed to join the closest major city) in addition to high daily temperatures and low average precipitations. Since lower temperatures and higher humidity values are usually associated with a longer virus survivability in the environment^38^, our results may suggest a more important role than what was previously believed of the indirect transmission through viral persistence in the environment for FMV in this region. Additionally, with the viral diffusion being positively affected by high cattle and human densities and negatively affected by large accessibility values, anthropological activities seem to have an impact on the virus diffusion. These observations could be the consequence of infected herds of cattle moving from smaller rural localities toward nearby larger cities with cattle markets^14,39^.

Regarding the effect of the different selected predictors on virus diffusion in a continuous setting, our results suggest that cattle densities above 125 cattle per km^2^ and the presence of cropland (pure cropland or mixed with other types of land) both have a negative impact on virus diffusion. Ours results suggest that the virus had difficulties to spread beyond the geographic region located at the root of the tree, where high cattle densities and low crop densities were present and to spread to areas with low cattle densities but high crop densities, presumably due to lack of suitable hosts. However, it is difficult to know exactly whether it is the cattle or crop density that had the most impact due to high correlation of the two variables at the time and in the region of origin.

The location uncertainty found at the root of the continuous tree could explain the differences between the discrete and continuous methods in estimating the effect of the cattle density on virus diffusion. For our analysis, this uncertainty seems to be translated by the *SEREPHIM* programme as a period where the virus is almost not moving. This uncertainty seems to drive *SEREPHIM* to the conclusion that the high cattle densities found near the origin of the epidemic are related to this lack of movement and therefore estimate that they have a negative influence on the virus diffusion. Although we suspect a link between the cattle density and the location of emergence of the analysed clade, we think that the continuous analysis does not offer the resolution needed to understand that relation (i.e. the spatial HPD confidence interval is too large). By parameterizing each rate of among-location movement as a function of predictors, the discrete approach seems therefore more appropriate to characterise the environmental and anthropological effect of the virus diffusion in this endemic situation.

In conclusion, the reconstituted phylogeographical tree pattern for the FMDV serotypes A, O and SAT2 reflects a situation where the recent FMDV circulation is mainly driven by commercial exchanges, through pastoral herd movements, and where wildlife seems to have almost no influence on the intra-continental circulation of the disease. However, the observed patterns for SAT2 reflects a situation where wildlife (wild buffaloes) constitute the original host of the serotype, whilst the observations for A and O suggest that those serotypes were imported in Africa at the start of the 19^th^ century. We observed that indirect transmission through the environment and direct transmission through anthropological activities had an enhancing effect on the virus diffusion in Eastern Africa.

## MATERIALS AND METHODS

### Data Collection

To obtain a comprehensive genetic dataset we first retrieved all available African FMDV A, FMDV O and FMDV SAT2 genetic sequences in Genbank (accessed on the 15/12/2016). From these datasets we selected all VP1 sequences with at least information on the country of sampling and the year of sampling. In total we gathered 191 FMDV A, 351 FMDV O and 477 FMDV SAT2 sequences. The sequences were aligned using Multiple Alignment Fast Fourier transformation (*MAFFT*) ^40^. Potential recombinant sequences were detected with RDP4 software and any such sequences were removed^41^.

To reduce the effect of potential sampling bias we removed all sequences coming from countries with less than three sequences available (in the whole-time span) and ran a stratified subsampling procedure to allow a maximum of three sequences per country of origin and per month. The spatial coordinates of sampling for each sequence was retrieved using the *GGMAP* package in R and the most precise sampling localisation name available for each sequence^42^. The final FMDV A dataset was composed of 107 sequences from eight countries, dates ranging from 1966 to 2016. The final FMDV O dataset was composed of 192 sequences from 12 countries, dates ranging from 1964 to 2016. The final FMD SAT2 dataset was composed of 135 sequences from 15 countries, dates ranging from 1970 to 2015 (see supp tables 15,16 and 17).

### Bayesian Evolutionary Inference

Discrete phylogeographic tree inference

Time-scaled phylogenetic trees were inferred using *BEAST* 1.8 with the *BEAGLE* library^43^, and different substitution clock and population evolution models were evaluated by estimating their marginal likelihoods using the Akaike’s Information Criterion for MCMC samples (AICM) in Tracer 1.6. Ultimately a general-time-reversible (GTR) model with site to site rate variation between two categories was selected as nucleotide substitution model^44^ with a Bayesian skygrid population model and a relaxed uncorrelated log-normal molecular clock model were chosen to model the evolution of the FMDA and FMDO serotypes. For the FMD SAT2 serotype a HKY nucleotide substitution model with a constant clock model with a Bayesian skygrid population model were chosen to model its evolution^45,46^. Posterior sets of trees were generated for each serotype by combining at least 2 independent Markov Chain Monte Carlo run of 40 million steps sampling every thousand with 10% burn-in.

We first reconstructed the time-scaled phylogenetic trees for the three studied serotypes. Thereafter to reduce the computation time of the GLM and the spatial diffusion analyses we estimated the spatial model components using subsets of 1000 trees from the original posterior distributions of trees as input empirical tree distributions. We used *TreeAnnotator* to summarize maximum clade credibility (MCC) trees and *FigTree* version 1.4.1 to visualize the annotated trees^47,48^. The software *SPREAD3* and *Cytoscape* were used to identify and visualize the well supported rates of transmission through a Bayes factor test ^49^.

For the three serotypes we reconstituted the discrete transition events between the different sampled countries through the whole phylogeny using the “mugration model”. Therefore, an asymmetric continuous-time Markov chain (CTMC) model with an incorporated Bayesian stochastic search variable selection (BSSVS) was used to determine which set of transition rates sufficiently summarizes the epidemiological connectivity between the countries^30^. A posterior inference of the complete Markov jump history through the whole genealogy was also performed, in order to quantify state transitions and the time spent in each country by the virus.

### Environmental and anthropological effect estimation

#### Monophyletic clade selection

Using the previously reconstructed discrete phylogreographic tree of the FMDV O serotype we selected a monophyletic clade with a MRCA under 25 years and a posterior probability over 50 % on the location for all its nodes. To avoid uncertainty in the predictor effect estimation analysis we removed all sequences connected to branches with length more than 10 years. At the end of the process the dataset was composed of 46 FMDV O sequences coming from 31 locations across Kenya, Uganda and Tanzania.

#### Generation of predictive factors of FMDV diffusion

A Generalised Linear Model (GLM) extension of the discrete approach was used to test and quantify the enhancing (positive) or impeding (negative) effect of potential predictors on the viral diffusion process^32^. This model parametrizes the transmission rate matrix between discrete locations as a log linear function of the potential predictive factor matrices. While reconstructing the phylogeographic history the model performs Bayesian model averaging to determine which combination of predictor matrices are the best to explain the spatial diffusion process. For each predictor a Bayes factor (BF) value is calculated based on the ratio of posterior to prior probabilities of inclusion ^50^.

The different predictors of FMD diffusion considered were: the accessibility to the sampled location, cattle density, crop density, the elevation of the location, the forest density, the human density, the average yearly precipitation, the shrubland area density, the average daily temperature (for the provenance see sup, table 7). Each potential predictor was retrieved as a raster matrix, representing the predictor spatial localisation, and aggregated to a resolution of 0.08 by 0.08, corresponding to pixels of approximately 8 km by 8 km.

The circuitscape software was used to determine the predictors values used in our GLM analysis^51^. For each predictor, two predictor values were generated, one using the raster as resistance values (impeding the viral diffusion) and the other using the raster as conductance value (enhancing the viral diffusion). To obtain those values we used a circuit theory approach to estimate modified distances, used as predictor values, between each pair of locations using the raster values as heterogeneity factors^50^. Consequently, if a raster was used as a resistance surface, we would estimate large predictor values between the locations separated by high raster values and small predictor values between the locations separated by small raster values. Prior to their inclusion in the GLM analyses the predictor values were log transformed and standardised. Each analysis was run by comparing the effect of a predictor with a null predictor, corresponding to a random raster.

Complementary to the discrete GLM approach, we tested and quantified the effect of the potential predictors using a continuous coordinate approach. Therefore, we inferred the diffusion of the virus using a random walk model of diffusion and used the SERAPHIM package to test and estimate the effect of the predictors on the virus diffusion^31,34^. Like the discrete approach, *SERAPHIM* estimates a modified distance for each pair of locations find at the start and end of the phylogeny branches. The correlation between the time spent on each branch and the estimated distance value is then estimated. The statistical significance of this correlation is tested using a randomized phylogeny and expressed in the form of a BF^34^.

## Acknowledgements

The study was supported by Roslin Institute Strategic grant funding from the the Biotechnology and Biological Sciences Research Council, UK (BBS/E/D/20002173). SL is supported by a Chancellor’s Fellowship from the University of Edinburgh, and FD is supported by a Chancellor’s Fellowship studentship.

## Author contributions

FD, SL and MB conceived the study; FD analysed and interpreted the data, and drafted the manuscript; SL and MB provided guidance on the study design and the analysis, and contributed to interpreting the data and scientific discussions. All authors read and approved the final manuscript.

## Additional information

Early pre-print of this manuscript on BioRxiv **doi:** https://doi.org/10.1101/358044 https://www.biorxiv.org/content/early/2018/06/28/358044.1

